# Sensory encoding of emotion conveyed by the face and visual context

**DOI:** 10.1101/2023.11.20.567556

**Authors:** Katherine Soderberg, Grace Jang, Philip Kragel

**Affiliations:** Emory University, Department of Psychology

## Abstract

Humans rapidly detect and interpret sensory signals that have emotional meaning. The posterior temporal sulcus (pSTS) and amygdala are known to be critical for this ability, but their precise contributions—whether specialized for facial features or sensory information more generally—remain contentious. Here we investigate how these structures process visual emotional cues using artificial neural networks (ANNs) to model fMRI signal acquired as participants view complex, naturalistic stimuli. Characterizing data from two archival studies (*N*s = 20, 45), we evaluated whether representations from ANNs optimized to recognize emotion from either facial expressions alone or the broader visual context differ in their ability to predict responses in human pSTS and amygdala. Across studies, we found that representations of facial expressions were more robustly encoded in pSTS compared to the amygdala, whereas representations related to visual context were encoded in both regions. These findings demonstrate how the pSTS operates on abstract representations of facial expressions such as ‘fear’ and ‘joy’ to a greater extent than the amygdala, which more strongly encodes the emotional significance of visual information more broadly, depending on the context.

## Introduction

Humans are constantly confronted with sensory signals that have emotional meaning. A grimace, slumped shoulders, the arrangement of people in a scene—all these sensory events convey affective information critical for navigating a dynamic environment. Though these signals are varied and complex, humans rapidly detect and interpret them. A particularly salient source of signals is facial expressions, which convey information about emotional states. Perceiving the emotion expressed by the face, humans can infer an individual’s intentions and motivations, which are crucial for successful social interaction. Along with facial expressions, other elements of the visual array such as actions, objects, and scenes can signify emotional meaning. Detecting the affective significance of one’s present environment powerfully motivates and constrains behavior. How does the brain accomplish this sensory encoding of emotion? Studies of face perception and emotion processing implicate the superior temporal sulcus (STS) and amygdala as important nodes in distributed neural networks for registering emotional information in the sensory environment.

Research in the field of cognitive neuroscience has revealed a distributed network of regions that process faces, extracting representations of who the face belongs to and the emotion it conveys. Bruce and Young’s influential model posited that cognitive processing of faces occurs in independent streams for identity and emotional expression^1^. Haxby and colleagues extended this idea, suggesting that facial features are processed by hierarchically organized systems for face processing^2^. In this and related models^3,4^, dorsal visual regions process changeable facial features and integrate them with other signals, while regions in the ventral visual stream process static features. The posterior STS is thought to integrate dynamic information from facial expressions, along with other social signals (e.g., body posture and vocal tone), and map expressions to emotion categories^5–7^. The amygdala, on the other hand, plays a role in extended face processing systems by sensing threat and other salient affective information from facial expressions and has been shown to discriminate between fearful and non-fearful faces^8–10^. Thus, the posterior STS and the amygdala have been robustly implicated in processing emotional faces, each with dissociable functions.

The posterior STS responds to a variety of sensory signals that guide the perception of facial expressions. Broadly, it is sensitive to biological motion, from minimal arrays to complex action sequences^11,12^. It is consistently engaged by facial expressions and its responses to these expressions are sufficient to decode emotion category^7,13^. Furthermore, the posterior STS responds to emotional cues originating from other sources, including tone of voice and body posture, which also contain information from which emotion category can be decoded^14,15^. This evidence implicates the posterior STS as crucial for processing the dynamic, multimodal nature of emotions expressed by others^16^, and has led some to suggest that the posterior STS encodes supramodal, or amodal representations of emotion^17^. Such representations, which consist of emotion concepts abstracted away from their inputs, result from the processing of modality-specific features from distinct sensory pathways that are mapped to more abstract categories.

The amygdala, a component of extended face processing systems, has been associated with the detection of salient exteroceptive events, including innate and learned threats^18,19^, rewards^20^, animate objects^21^, and multiple different facial expressions^22^. Evidence is mixed as to whether the amygdala responds preferentially to one or several facial expressions as opposed to facial emotion more generally. A long line of studies shows that the amygdala preferentially responds to fearful (and sometimes angry and disgusted) expressions compared to neutral ones^23–26^. More recent work using multivariate decoding approaches found that amygdala activity could differentiate fearful from non-fearful faces, but not other categories^10^. However, there is evidence against specificity for fear, as amygdala activity has been found to reflect the intensity or ambiguity of facial expressions rather than a single category such as fear^27,28^. Given its broader role in processing both threats and rewards, amygdala responses to facial expressions of fear and anger have most commonly been interpreted as involving the detection of threat^29^ or salience more broadly^30–32^. Recent evidence suggests that the amygdala does not always preferentially respond to facial emotion compared to non-faces^33^. Debate continues over whether amygdala responses to expressions of fear reflect holistic encoding of that emotion category, or whether this activity reflects representations related to broader dimensions of salience, valence, or intensity rather than representations of facial expressions themselves.

Given this evidence, it is clear that the amygdala and posterior STS are involved in processing emotional facial expressions. However, much of our understanding of these two regions is based on research using paradigms with decontextualized facial expressions, such that the face dominates the visual array. For example, a prominent paradigm shows subjects cropped fearful and angry faces, devoid of any context^34^. In this and similar tasks, observing greater activity in the amygdala or posterior STS could mean that these regions encode representations of specific facial expressions, or that they encode representations related to the broader visual context that are not specific to facial expressions *per se*. Accordingly, it is an open question as to the generalizability of representations encoded in these regions; it is possible that the information encoded by either or both of these regions is better explained by visual context as opposed to facial expressions specifically. Examining social emotion processing in more naturalistic situations enables tests of generalizability; however, increased ecological validity lowers experimental control and makes it challenging to parse which elements of a multifaceted stimulus drive brain responses.

Here we address this problem by evaluating the nature of representations encoded in the amygdala and posterior STS in a naturalistic context using artificial neural networks (ANNs) as models of human brain systems. ANNs are trained to perform a particular task, and through this process they learn representations useful for a specific computational goal, such as object and face recognition^35^. Of the many possible ways a task can be accomplished, ANNs with the right set of inductive biases will be able to efficiently achieve high performance^36^. From a design optimization perspective, it been argued that the representations learned by task-optimized ANNs are fundamentally related to those learned by biological neural networks because both systems solve the same tasks with similar inductive biases^37,38^. Therefore, we use the term “representation” to refer to patterns of activity in both biological and artificial neural networks, given the hypothesized link between them.

Modeling efforts using ANNs to approximate the ventral visual stream have provided multiple insights about object recognition^39^ and face processing^40–42^, including functional specialization in the fusiform gyrus^43^. However, they have been less successful at explaining activity in dorsal and lateral pathways^44^, and emotion recognition in particular. Further, despite the ability of ANNs to parse naturalistic features from a complex, multimodal stimulus, this work has largely characterized brain responses to tightly controlled experimental paradigms that do not mimic experiences from everyday life because they do not present faces in rich social contexts with concurrent multimodal stimuli^45^. We address these issues by testing whether ANNs with different objectives can better explain the functions of the posterior STS and amygdala to naturalistic videos. We compare the performance of a deep convolutional network with attention mechanisms trained to recognize facial expressions and predict the location of specific facial landmarks (EmoFAN)^46^ and a deep convolutional network trained to classify the emotion schema present in visual scenes (EmoNet)^47^. These models have been trained and initially validated on a range of standard facial expression and naturalistic scene databases^46–48^, showing that they use distinct representations to classify emotional signals (Supplementary Text and Supplementary Figures 1-3). Using features extracted by these ANNs to develop encoding models that predict brain activity in a given region^49^, we characterize how the amygdala and posterior STS disentangle the emotional meaning of facial expressions and the broader visual context.

## Results

### Functional dissociation of emotion processing

To investigate face processing in a rich naturalistic context, we examined human brain activity in two studies. In Study 1, subjects (*N* = 20) watched the full-length movie *500 Days of Summer* while undergoing fMRI (∼1.5 hours per participant; data sampled from the Naturalistic Neuroimaging Database^50^). Using the deep convolutional neural networks trained for facial and scene classification, we extracted features from each timepoint in the movie and fit encoding models to predict brain activity in the amygdala and posterior STS (Figure 1). This enabled us to test how two models of emotion processing with different architectures, training objectives (expression-specific vs. general visual context), and sensitivities to different emotional signals predict the functional response of each region.

**Figure 1.**
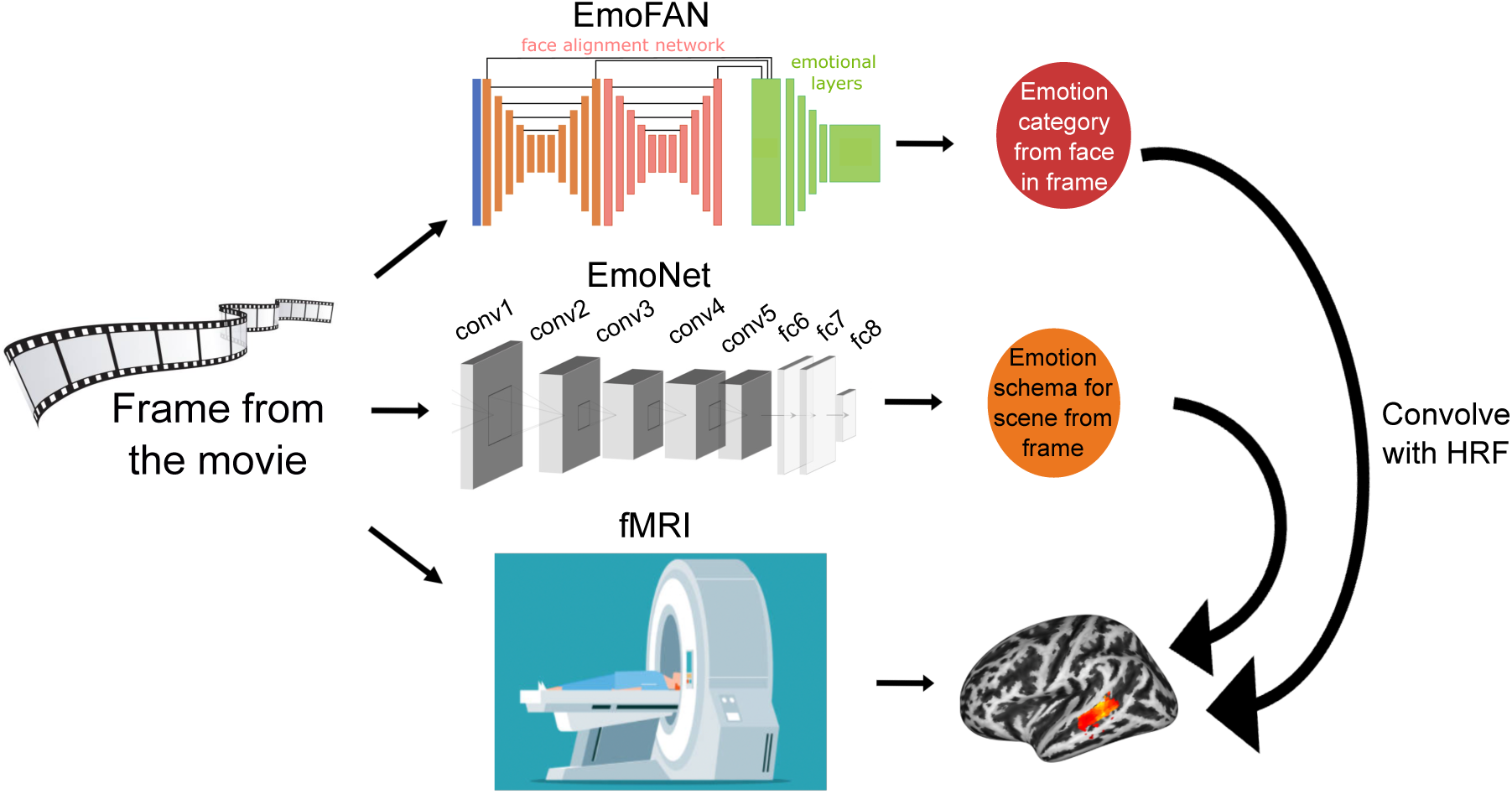
Encoding models for facial expressions and visual context. Every fifth frame of the movie is fed into two artificial neural networks. The upper ANN (EmoFAN) identifies emotional facial expressions, while the lower ANN (EmoNet) classifies the emotion schema of the image. Activations from layers of each of these ANNs are convolved with the hemodynamic response function (HRF) to create an encoding model of predicted brain activity based on facial expression features for EmoFAN and features related to visual context for EmoNet. These variables are used as features in multivariate encoding models to predict BOLD activation in the brains of subjects watching the full-length movie.

Given evidence of the distinct contributions of the amygdala and posterior STS to face processing, we hypothesized that posterior STS encodes representations related to facial expressions, and that the amygdala encodes representations related to the emotional significance of visual context more broadly. If this is the case, we would expect a double dissociation, such that abstract representations of facial expressions from EmoFAN would meaningfully predict activity in posterior STS and not the amygdala, with the opposite pattern for abstract emotion representations in EmoNet. However, given the evidence that the posterior STS responds to visual signals of emotion and may contain representations of emotion over and above those related to specific facial expressions, visual context (captured by EmoNet) might alternatively be encoded in posterior STS as well as the amygdala. In addition, we hypothesized that along with abstract emotion representations, more fine-grained features (such as raised eyebrows or the sclera of the eyes) might be also encoded in these regions.

To evaluate these alternative accounts, we developed multivariate encoding models using features from the final layer of each ANN; for EmoFAN, this consisted of a 10-dimensional layer capturing the probabilities of 8 emotion categories as well as continuous variables for valence and arousal; for EmoNet, this consisted of a 20-dimensional layer of 20 emotion categories (for more detail on network architectures, see Methods). Additionally, to capture low-level features, we developed models using intermediate layers of each ANN, which are ultimately combined to compute the final objective of each network.

We first evaluated whether features related to facial expressions are encoded in posterior STS, with the goal of testing whether emotional information from multiple sources are jointly encoded in this region. We found evidence that both encoding models—the one based on facial expressions and the one based on visual context—explained activity in posterior STS (average EmoFAN prediction-outcome correlation = .054, *SD* = .012, 95% CI = [.049 to .060], Cohen’s *d* = 4.42, 67.8% noise ceiling; average EmoNet prediction-outcome correlation = .086, *SD* = .018, 95% CI = [.079 to .095], Cohen’s *d* = 4.67, 70.5% noise ceiling; Figure 2a-b). Voxelwise inference (one-sample t-test, FDR *q* < .05, two-tailed) revealed that a broad extent of voxels in bilateral posterior STS were predicted by both EmoFAN (peak effect in dorsal left posterior STS: extent = 285 voxels/9328 mm^3^, *t* = 13.2, location in MNI space = [-56, -35, -2]; peak effect in ventral right posterior STS: extent = 357 voxels/11680 mm^3^, *t* = 18.28, location in MNI space = [56, -29, -5]) and EmoNet (peak effect in dorsal left posterior STS: extent = 283 voxels/9264 mm^3^, *t* = 16.94, location in MNI space = [-56, -35, -2]; peak effect in ventral right posterior STS: extent = 361 voxels/11824 mm^3^, *t* = 15.71, location in MNI space = [56, -29, -5]).

**Figure 2.**
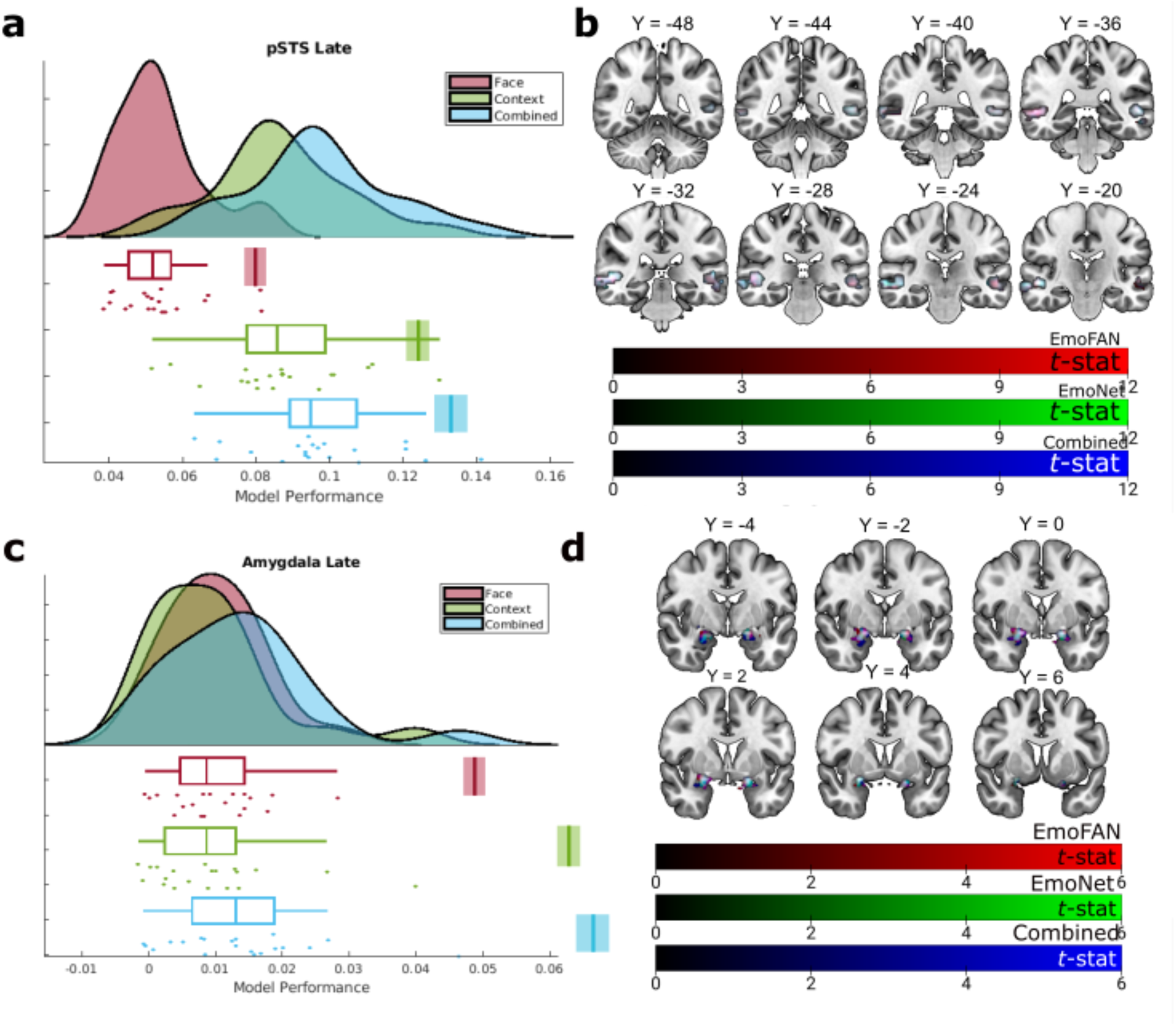
Encoding of abstract facial emotion and emotion schemas in the posterior STS and amygdala. (**a, c**) Encoding model performance using features from the late layers of artificial neural networks predict posterior STS and amygdala activity. EmoFAN shown in maroon, EmoNet in green, and the combined model in blue. Density and boxplots show model performance across subjects; each dot represents that model’s performance on one subject’s brain data (*N* = 20). Estimated noise ceilings determined by resubstitution are indicated by the vertical bars (standard error in lighter shade). (**b, d**) Voxels predicted by EmoFAN, EmoNet, and the combined model are overlapping throughout the posterior STS and amygdala, thresholded at *q*_FDR_ < .05. Overlays show encoding performance against chance, with EmoFAN in red, EmoNet in green, and the combined model in blue.

Because the amygdala has been implicated in rapid processing of certain facial features, particularly those related to rapid orienting towards threat-relevant emotions like anger and fear^28,51^ rather than multiple abstract categories, we hypothesized that abstract representations in later layers of EmoFAN would not be robustly encoded in patterns of amygdala activity. Inconsistent with our hypothesis, both encoding models predicted response patterns in the amygdala (average EmoNet prediction-outcome correlation = .009, *SD* = .010, 95% CI = [.006, .015], Cohen’s *d* = .949, 13.3% noise ceiling; average EmoFAN prediction-outcome correlation = .010, *SD* = .007, 95% CI = [.007, .014], Cohen’s *d* = 1.41, 20.1% noise ceiling; Figure 2c-d).

Voxelwise inference revealed that activity in bilateral basolateral amygdalae were predicted by EmoNet (Figure 3b; peak effect in right amygdala: extent = 87 voxels/2848 mm^3^, *t* = 7.65, location in MNI space = [23, -2, -17]; peak effect in left amygdala: extent = 70 voxels/2288 mm^3^, *t* = 7.05, location in MNI space = [-20, -2, -17]) and EmoFAN (peak effect in right amygdalostriatal area: extent = 8 voxels/264 mm^3^, *t* = 4.11, location in MNI space = [29, -11, -11]; peak effect in right basolateral amygdala: extent = 110 voxels/3600 mm^3^, *t* = 7.74, location in MNI space = [23, -2, -17]; peak effect in left basolateral amygdala: extent = 87 voxels/2856 mm^3^, *t* = 8.05, location in MNI space = [-20, -2, -17]).

**Figure 3.**
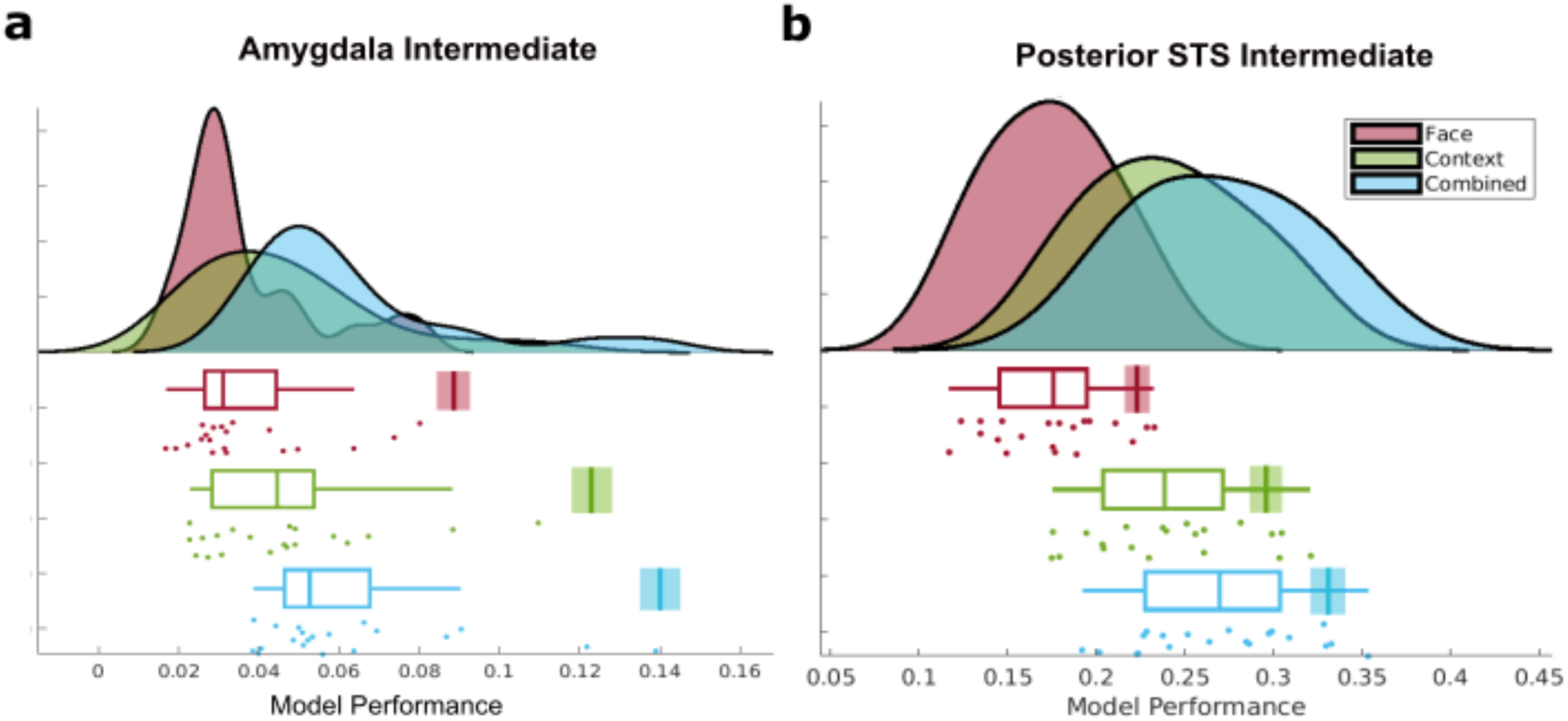
Intermediate layers of both encoding models, as well as the combined model, predict activity in the amygdala and posterior STS. Performance of encoding models using features from the intermediate layers of each model predicting activity in the (**a**) amygdala and (**b**) posterior STS (*N* = 20). EmoFAN is shown in maroon, EmoNet is shown in green, and the combined model is shown in blue. Density and boxplots show model performance across subjects; each dot represents that model’s performance on one subject’s brain data. Estimated noise ceilings determined by resubstitution are indicated by the vertical bars (standard error in lighter shade).

To determine whether representations of facial expressions and visual context each explain unique variance in posterior STS and amygdala activity, we created a joint encoding model for each region that combines features from both ANNs. In this comparison, if the representations learned by the two ANNs are redundant, the combined model should do no better than either individual model, whereas if each model contains distinct information, the combined model should have a higher predictive value. Although both models predicted activity in each region, this comparison provides a strong test of our hypothesis that facial expressions are encoded more robustly in posterior STS compared to amygdala.

We compared encoding model performance using a 2-way ANOVA and found a region by model interaction (F_2,38_ = 112.69, *p* < .001, partial η^2^ = .856). The joint encoding model explained more posterior STS activation than either model alone (average prediction-outcome correlation = .097, *SD* = .019, 95% CI = [.089, .016], Cohen’s *d* = 4.97, 71.9% noise ceiling; difference between single (EmoNet) and combined = .011, *SD* = .004*, p* < .001, CI = [.008, .013]), suggesting that both types of representations are uniquely encoded in posterior STS. In contrast, there was a much smaller difference between the combined model and the visual context model alone in the amygdala (average prediction-outcome correlation = .014, *SD* = .011, 95% CI = [.010, .019], Cohen’s *d* = 1.24, 17.9% noise ceiling; difference between single (EmoNet) and combined = .004, *SD* = .004, *p* < .001, CI = [.002, .007]), suggesting that abstract representations of facial expressions predict a substantial amount, but comparably less unique variance in amygdala activity.

To determine which voxels in posterior STS were better predicted by the combined model compared to either EmoFAN or EmoNet alone, we performed group inference across subjects and found that voxels in bilateral posterior STS were better predicted by the combined model compared to either model alone (Supplementary Figure 2; peak effect in left dorsal posterior STS: extent = 285 voxels/9336 mm^3^, *t* = 21.61, location in MNI space = [-56, -35, -2]; peak effect in right ventral posterior STS: extent = 361 voxels/11824 mm^3^, *t* = 20.01, location in MNI space = [56, -29, -5]).

Similarly, to determine which voxels in the amygdala were better predicted by the combined model compared to either EmoFAN or EmoNet alone, we performed group inference across subjects and found that voxels in bilateral amygdala were better predicted by the combined model compared to either model alone (Supplementary Figure 3; peak effect in right basolateral amygdala: extent = 151 voxels/4960 mm^3^, *t* = 8.45, location in MNI space = [23, -2, -17]; peak effect in left basolateral amygdala: extent = 113 voxels/3688 mm^3^, *t* = 6.69, location in MNI space = [-20, -2, -17]; peak effect in left amygdalostriatal transition area: extent = 7 voxels/216 mm^3^, *t* = 2.78, location in MNI space = [-29, -5, -11]).

Because the amygdala and posterior STS are known to respond to low level visual features present in faces, including the high contrast present in the eyes and low-frequency information about face shape^11,52,53^, it is possible that activity in earlier layers of both ANNs— which are ultimately transformed into representations of emotion categories—might predict responses in these areas. To examine the predictive ability of encoding models based on these more basic features, we fit a new set of encoding models using features from earlier layers. We found that both of these encoding models explained activation in the amygdala (Figure 3; average EmoFAN prediction-outcome correlation = .037, *SD* = .018, 95% CI = [.031 to .046], Cohen’s *d* = 2.10, 40.6% noise ceiling; average EmoNet prediction-outcome correlation = .046, *SD* = .022, 95% CI = [.039 to .058], Cohen’s *d* = 2.05, 36.0% noise ceiling) and posterior STS (average EmoFAN prediction-outcome correlation = .172, *SD* = .034, 95% CI = [.158 to .186], Cohen’s *d* = 5.01, 80.1% noise ceiling; average EmoNet prediction-outcome correlation = .235, *SD* = .042, 95% CI = [.218 to .254], Cohen’s *d* = 5.35, 80.0% noise ceiling). These results demonstrate that more basic visual features related to emotional expressions and visual context explain activity in both regions.

To assess the dependence of encoding model performance on depth (intermediate vs. late layer), region (amygdala vs. posterior STS), and model (EmoFAN, EmoNet, and combined), we performed a 3-way ANOVA and found a region by depth by model interaction (*F*_2,38_ = 54.925, *p* < .001, partial η^2^ = .743). A planned contrast revealed this interaction was driven by a greater performance for pSTS encoding models (relative to the amygdala) fit using features from EmoFAN (compared to EmoNet) using late layers (compared to earlier layers; *t*_19_ = 6.06; *p* < .001). To examine the effect of feature type for models based on intermediate layers, we performed a follow-up 2-way ANOVA and found a region by model interaction for intermediate layers (*F*_2,38_ = 310.33, *p* < .001, partial η^2^ = .942). This interaction was driven by a larger effect of model type in posterior STS, such that the combined model showed a stronger additive effect (i.e., more variance explained by features from EmoFAN) in posterior STS compared to the amygdala. Overall, these results demonstrate that abstract representations of facial emotions are less predictive of amygdala responses to naturalistic videos compared to posterior STS.

Given functional imaging and neuropsychological research demonstrating right hemisphere dominance in face^54,55^ and emotion processing^56,57^, we additionally conducted an exploratory *post hoc* analysis comparing the performance of encoding models between hemispheres for both the pSTS and amygdala. We found minimal differences in encoding model performance between hemispheres for all models and regions (although some effects in the amygdala exhibited trends toward left hemisphere lateralization; Supplementary Table 4), indicating that representations learned by the ANNs were not encoded in lateralized brain systems.

### Positioning pSTS and amygdala in face processing networks

The posterior STS and amygdala contribute to face perception through their participation in a distributed neural network. To probe this network and estimate how representations of emotion from facial expression and context are encoded in other face-processing regions, we developed encoding models that predict activity in regions of cortex likely to contain the fusiform face area (FFA) and occipital face area (OFA)^58^. We selected these regions due to their roles in the core face processing system^2–4^, as they operate on static facial features (holistic configurations and parts of faces, respectively) and serve as inputs to the pSTS and amygdala. If it is the case that activity in these early face regions solely reflects feedforward processing of static features, then representations of ANNs for emotion categorization should poorly predict activity in either region. However, recent findings raise the possibility that these areas may not be functionally independent of downstream regions, as decoding studies show that FFA^7,59,60^ and OFA^60^ contain information about facial expression. Additional evidence indicates that OFA and FFA are functionally coupled with the pSTS and amygdala^4,61,62^, suggesting that information about emotion categories may be encoded throughout core and extended face systems.

Encoding models predicted activity in OFA and FFA with a similar performance as posterior STS activity (Supplementary Table 6). This suggests that features related to facial expression categories are encoded in regions throughout the distributed face processing network. However, while model performance was quantitatively similar across posterior STS, FFA, and OFA, this does not require that the same variables are encoded in each region. To assess whether the models capture distinct information conveyed in each region despite exhibiting similar levels of performance, we performed a similarity analysis on voxelwise model outputs for the late layer of EmoFAN. Predicted responses in the FFA and OFA were moderately similar (*r* = .247, *sem* = .026, Cohen’s *d* = 2.04), whereas correlations between posterior STS and either of these regions were weak (STS-FFA: *r* = .0399, *sem* = .025, Cohen’s *d* = .360, STS-OFA: *r* = .0400, *sem* = .024, Cohen’s *d* = .377; Figure 4). Correlations between core face regions and the amygdala exhibited intermediate correlations (amygdala-FFA: *r* = .1148, *sem* = .020, Cohen’s *d* = 1.26, amygdala-OFA: *r* = .0896, *sem* = .019, Cohen’s *d* =1.04). Furthermore, the correlation of predicted values was much stronger between voxels in OFA and FFA than between all other region pairs (*t*_19_ = 6.77, *p* < .001). These observations suggest that despite exhibiting similar performance, the nature of representations differs between regions implicated in processing invariant features of faces and posterior STS, a region thought to process dynamic expression features.

**Figure 4.**
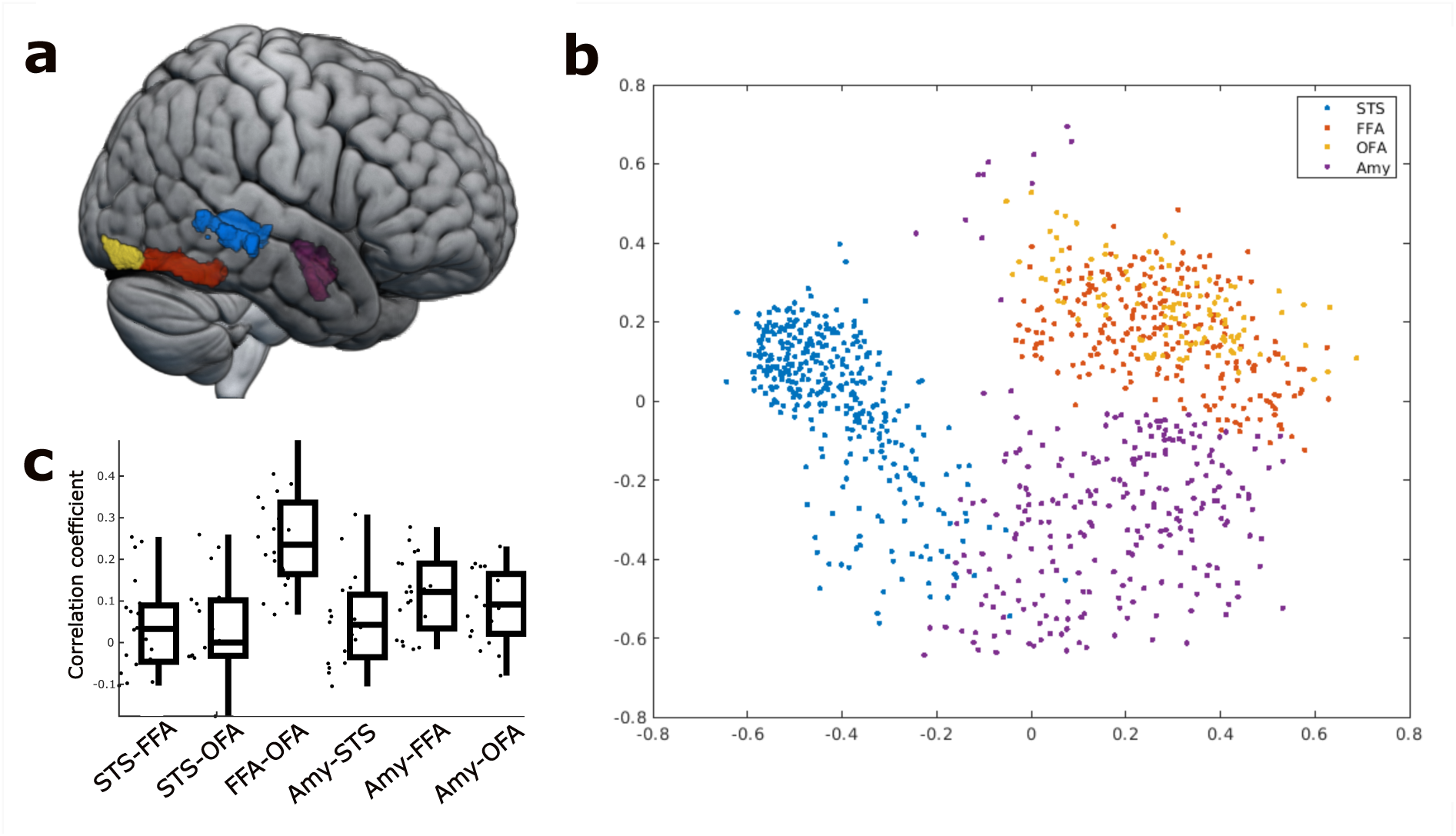
Encoding of abstract facial emotions in static face regions are much more similar than relationships between other regions involved in face perception. (**a**) Occipital face area (OFA, yellow), fusiform face area (FFA), posterior superior temporal sulcus (pSTS, yellow) and amygdala (Amy, purple) are rendered on the IBM152 template. (**b**) Model predictions visualized with multidimensional scaling performed on group average data. Each point corresponds to voxel-wise encoding model predictions. (**c**) Similarity of encoding model predictions between different pairs of regions of interest. Error bars reflect the standard error of the mean.

### Conceptual replication in a more controlled, yet naturalistic dataset

Study 1 revealed that both posterior STS and amygdala encode representations of facial expression and visual context, and that representations of facial expressions predicted more unique variance in posterior STS compared to the amygdala. Multiple factors could have contributed to these results. Scanner hardware (1.5 T field strength), pulse sequence, and the nature of the stimulus (a full-length motion picture with narrative elements) could have limited the performance of encoding models. These factors might have led to a reduced ability to detect signal in the amygdala, though the fact that amygdala was predicted by intermediate and late layers of each ANN lessens this concern. To examine the generalizability of the findings from Study 1, we performed a conceptual replication using archival data from an independent experiment that varied along these factors (Study 2).

Study 2 investigated face processing in 45 subjects as they watched a compilation of short clips taken from the television series “Game of Thrones” with concurrent fMRI^63^. As this stimulus involves decreased contextual variability, fewer narrative components, and an increased focus on the face compared to the film used in Study 1, it comprises a more constrained test of neural encoding. As a result, encoding models fit to data in Study 2 should be more sensitive to face processing and less reflective of naturalistic processing involving narrative and a wider range of contexts. Following the same procedures used to model brain responses acquired in Study 1, we used the features extracted by each ANN in encoding models to predict activity in the amygdala and posterior STS.

Replicating the findings from Study 1, we found that late layers from both models predicted activity in posterior STS (average EmoFAN prediction-outcome correlation = .266, *SD* = .044, 95% CI = [.254, .279], Cohen’s *d* = 5.77, 53.7% noise ceiling; average EmoNet prediction-outcome correlation = .401, *SD* = .063, 95% CI = [.382, .419], Cohen’s *d* = 5.66, 80.6% noise ceiling; Figure 5a). Additionally, we found that the combined model outperformed each individual model (average prediction-outcome correlation = .433, *SD* = .065, 95% CI = [.414, .451], Cohen’s *d* = 5.80, 80.9% noise ceiling), suggesting that representations of facial expressions and visual context are encoded in this region across varying naturalistic contexts.

**Figure 5.**
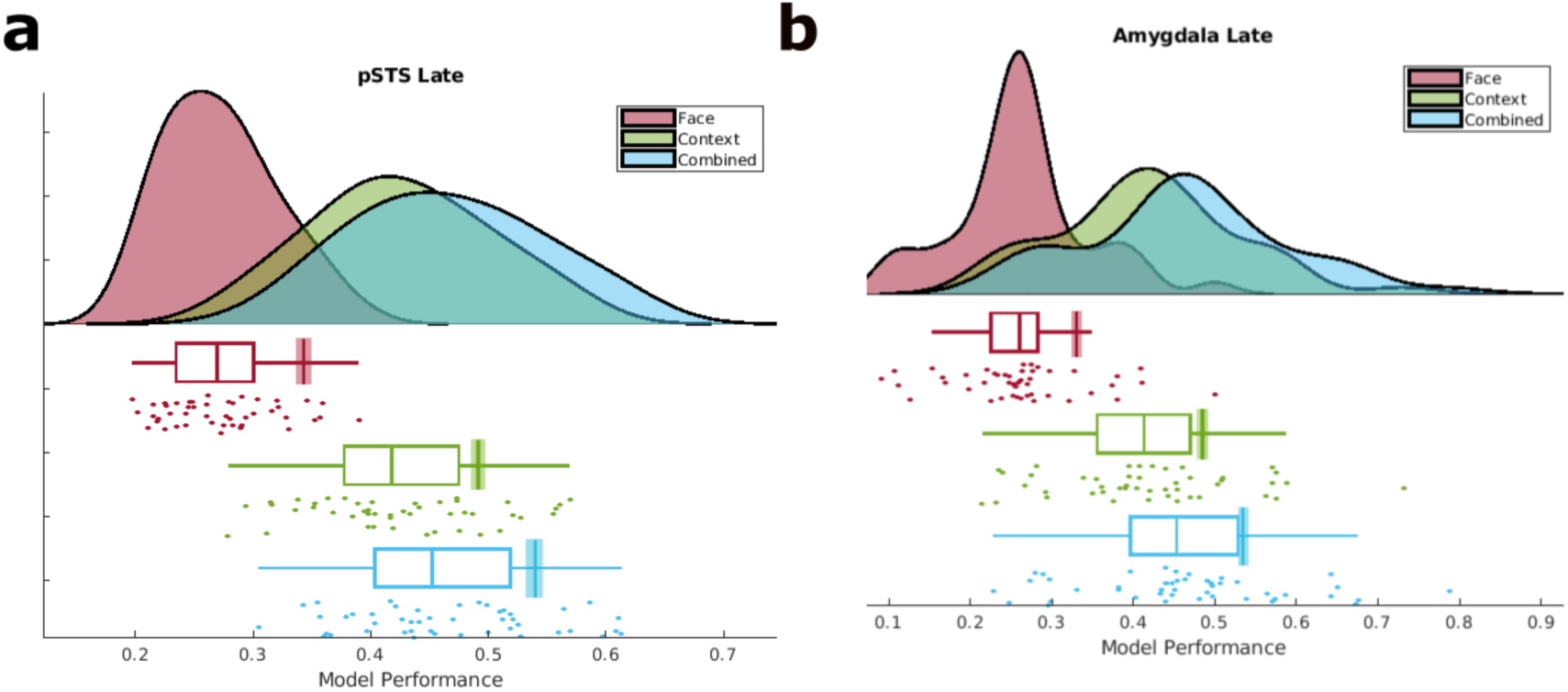
Late layers of both encoding models, as well as the combined model, predict activity in posterior STS and amygdala (Study 2). EmoFAN is shown in maroon, EmoNet is shown in green, and the combined model is shown in blue. Density and boxplots show model performance across subjects (*N* = 45); each dot represents that model’s performance on one subject’s brain data. Estimated noise ceilings determined by resubstitution are indicated by the vertical bars (standard error in lighter shades).

Additionally replicating the results of Study 1, we found that late layers of both models predicted amygdala activity (average EmoFAN prediction-outcome correlation = .251, *SD* = .075, 95% CI = [.230, .274], Cohen’s *d* = 3.20, 74.7% noise ceiling; average EmoNet prediction-outcome correlation = .388, *SD* = .090, 95% CI = [.362, .414], Cohen’s *d* = 3.80, 78.6% noise ceiling; Figure 5b). In addition, we found that the combined model outperformed both individual models (average prediction-outcome correlation = .425, *SD* = .098, 95% CI = [.394, .452], Cohen’s *d* = 3.76, 79.6% noise ceiling). However, in contrast to Study 1, we found no region by model interaction for late layers (F_2,88_ = .648, *p* = .525, partial η^2^ = .015), suggesting that abstract facial expression and visual context information did not substantially differ across regions.

We again found a region by depth by model interaction, replicating the results of Study 1 (*F*_2,88_ = 19.6, *p* < .001, partial η^2^ = .308; For model performance in intermediate layers, see Supplementary Table 7). As in Study 1, this interaction was driven by a difference in the predictive performance of encoding models, such that later layers (compared to earlier layers) of EmoNet (compared to EmoFAN) performed relatively better at predicting amygdala activity (compared to posterior STS). This suggests that as fine-grained visual context representations are transformed into more abstract representations, they predict amygdala activity to a greater extent than posterior STS.

## Discussion

This work provides a more precise understanding of how visual information about facial expressions and emotional context is encoded in the brain. Across two studies, we found that posterior STS encodes abstract emotion representations related to facial expressions. In data that examined responses to a contextually variable stimulus (Study 1), we found that abstract facial expression information was less predictive of amygdala activity compared to posterior STS; however, when predicting responses to a more constrained stimulus (Study 2) we found similar magnitudes of prediction across regions. In addition, across both studies we found that abstract emotion representations related to visual context are encoded in both amygdala and posterior STS. Furthermore, in both studies an encoding model combining features related to both facial expression and visual context explained a greater degree of posterior STS and amygdala activation than either model alone. These findings indicate that across contexts, the posterior STS processes visual information related to emotion from both facial expressions and the broader visual context, whereas the amygdala more robustly represents visual context and encodes specific categories of facial expressions in a more context-dependent manner. More broadly, this work suggests that ANNs with varying objectives, including face localization, expression recognition, and schema classification can explain activity in brain regions underlying social perception.

Our findings inform cognitive models of face perception: in particular, how the amygdala, posterior STS, FFA, and OFA contribute to the distributed network of brain regions that processes faces. We found that amygdala representations of facial expressions were robustly engaged when subjects viewed a series of clips from a television show depicting brief episodes with no clear narrative links and were weakly engaged as subjects watched a full-length movie with narrative structure and varied visual scenes. This pattern of results suggests that amygdala representations of facial expressions depend on context, dovetailing with findings from controlled experiments showing that amygdala responses are modulated by affective context and expectations^64–68^. Furthermore, encoding models using lower-level visual features consistently predicted amygdala activity across studies, demonstrating that the amygdala responds to more low-level visual representations that could be used to classify facial expressions. These observations are consistent with evidence suggesting that the amygdala receives information about low spatial frequency components of faces to orient to the eye region and guide behavioral responses to threat and reward^31,52,53,69^. Taken together, the performance of amygdala encoding models suggest that cognitive accounts of face processing should consider how the prioritization of visual information depends on temporal context.

Our results additionally shed light on the role of posterior STS in face processing. Debate continues over which set of variables are encoded in this region—whether it is specialized for processing facial movement or whether it takes modality-specific information as input and maps it to supramodal representations of emotion categories^17,70^. Across two studies we found that abstract emotion features derived from both facial expressions and visual context predict unique components of posterior STS activity; that is, the joint encoding model based on late layers from both EmoFAN and EmoNet predicted a greater proportion of posterior STS activity than either model alone. If posterior STS were only encoding abstract facial emotion categories, we would expect the combined model to perform no better than the single models. We thus conclude that it is unlikely that the only function of posterior STS is to represent abstract emotion categories that have been completely disentangled from their sensory source, as some have argued^14,17^. We further found that intermediate and late layers of both the face and the context model predicted posterior STS activity during naturalistic movie viewing. This suggests that posterior STS encodes both types of representations: those that are fine-grained as well as those that are more abstract and tied to emotion categories. Likely, multiple neural populations within these regions represent emotion signals at differing levels of granularity.

Our results also inform accounts of the FFA and OFA, which comprise earlier stages of face processing. Encoding model predictions for FFA and OFA were more similar to one another than to posterior STS and amygdala, a finding consistent with the idea that these areas represent stable facial features that do not vary over time^61^. Additionally, we found that abstract facial expression categories from EmoFAN are encoded in all regions, which adds to the existing evidence showing that information relevant to facial expression is processed in many regions^59,60^. While an exhaustive comparison between ANNs trained to recognize facial expression and identity is outside the scope of the current investigation, our results are in general agreement with a recent proposal that there are parallel streams involved in face identification and the retrieval of emotion concepts^62^.

This study demonstrates the utility of using ANNs as models of perceptual systems beyond object recognition in the ventral visual stream, and in particular shows that comparing ANNs with varying objectives is a useful tool for testing theory. Several studies have examined how well deep convolutional networks optimized for object recognition (e.g., AlexNet) explain activity in posterior STS^7172^. One study found that representations from an intermediate layer of a convolutional network predicts pSTS responses distinctly from hand-coded features related to perceived affect and social interaction^71^. The current work is consistent with these findings, in that intermediate features of EmoNet, which was finetuned from AlexNet, predicted pSTS responses. Moreover, a recent study found that late layers of deep convolutional networks trained to individuate static images of faces are poor predictors of dynamic face processing^72^. Here we find that an ANN trained specifically for facial expression categorization can meaningfully predict brain activity in posterior STS.

Crucially, our work extends the ANN-derived encoding model approach by using ANNs trained for specific objectives that match the putative tasks of the brain regions of interest. Examining the mappings between ANNs and brain activity allows for a finer-grained understanding of the neural computations involved in transforming visual input into representations that could be used to recognize emotion in the face. The ANNs we examined here have different objectives and architectures, with EmoFAN standing apart from most deep convolutional networks including those that categorize faces^73^ by including an initial filter that detects the current pose of a face and identifies important landmarks, which then influence a set of convolutional layers that operate on facial features at progressively lower dimensionalities. With this relatively straightforward architecture, we see that an EmoFAN-derived encoding model predicts posterior STS activity. Although the amount of variance explained is far from characterizing all activity in this region, and the structure of EmoFAN is far simpler than that of the face processing systems in humans, the representations that EmoFAN has learned are meaningfully related to human brain activity, and thus advance our understanding of social information processing in the brain.

EmoFAN is trained on static images of facial expressions; given the inherently dynamic, multimodal nature of social emotion perception^74^, there are clear ways we could improve prediction using the ANN framework. To better capture the fact that posterior STS processes emotion signals from multiple modalities, we could incorporate features from ANNs that process auditory and linguistic inputs. This would allow us to test theories about whether and how posterior STS integrates multimodal inputs into a supramodal representation. Additionally, given evidence that posterior STS processes changing visual features, ANNs that can represent dynamic sequences of facial expressions would likely improve prediction. For instance, applying a long short-term memory architecture^75^ to representations from EmoFAN and other face-processing ANNs is a promising possibility in this vein^76^. To capture the complex process of social emotion perception more accurately, future work should explore how including features from other modalities, as well as accounting for dynamic sequences of expressions, might improve explanatory power. Additionally, establishing associations between ANNs and neural processing of emotion signals lays the groundwork for causal testing. For instance, ANNs with varied inductive biases can inform theories of cognitive development and functional specialization^42^; their features can also be used as inputs for stimulus generation designed to target particular neural populations.

This work is among a number of studies employing naturalistic paradigms to study social perception^71,77,78^. Naturalistic approaches allow us to examine how the brain represents the multifaceted, complex nature of emotion; however, it is challenging to analyze experiments of this type. Stimuli are not presented in a controlled manner, which makes it difficult to isolate the effects of any one factor or component. Our approach, in which ANNs are used to extract complex features related to emotion, allows multifaceted stimuli to be parsed and corresponding neural responses to be probed. In this framework, any neural network can be used to extract features related to the task it has been trained for, and these features can be tested for the degree to which they map on to brain activity^79^. Importantly, this allows us to better understand the brain as it is functioning in a more ecologically valid context. In the real world, it is rare that humans perceive others’ emotions in a vacuum, and we can better capture such contextually embedded processes using the present approach.

The ability of ANNs to perform a complex social behavior and predict human brain function highlights fruitful directions for future research. Although EmoFAN and EmoNet can accomplish the tasks they were trained for, they each constitute only one way of solving the given problem that stems from how they were trained; there exists a wide space of neural networks that accomplish similar tasks. In addition, subjects in both studies were not tasked with identifying the emotions from the faces they saw, and while emotional faces are likely attended to due to their salience^80^, we did not evaluate brain responses to face processing during an explicit recognition task. Additionally, the study samples were designed to characterize within-subject effects (*N*s = 20, 45) and are thus unable to robustly characterize any sex-related, cultural, developmental, or racial impacts on how the brain processes emotional faces. In addition, our approach does not causally test how information is conveyed between the amygdala, posterior STS, and other regions in the face processing network. Some work has found evidence that they are causally connected by showing that transcranial magnetic stimulation applied over STS during face perception leads to a reduced amygdala response^81^. Here we do not test whether information is transmitted between these regions.

The overall proportion of brain activity explained by encoding models in the pSTS and amygdala was small, likely due to the constrained nature of each ANN and the complexity of the naturalistic free-viewing paradigms. This illustrates how detecting emotion conveyed by visual signals is but one of many functions performed by these regions. Past work has used tightly controlled experimental paradigms that aim to minimize context effects and variation in brain responses over time^39,82^. Experiments of this kind minimize noise, but they may miss out on meaningful aspects of brain function, including multimodal processing and integration that occur in more ecologically valid paradigms. It is likely that more complex models will be necessary to fully explain the function of these brain regions during emotion perception. Future modeling efforts could focus on the relationships between these and other regions to further illuminate the distributed brain network involved in processing emotional visual information.

By testing ANN-derived encoding models on brain activity during naturalistic movie viewing, we have shed light on how signals of emotion are represented in the brain. The emotional meaning of the environment can be sensed from multiple sources, including others’ facial expressions and visual cues related to emotional context. Our findings suggest that the amygdala represents emotional context more robustly than specific facial expressions, whereas the posterior STS operates on both facial expression-specific representations and abstractions from sensory inputs to emotion categories. Further exploration using neural network-derived encoding models will deepen our knowledge—not only *that* emotion signals are processed in these regions, but *how* they convert visual inputs into an abstract representation of the present emotion state. Doing so will increase our understanding of the neural computations that enable the human brain to take in the complex emotional world.

## Methods

### fMRI Data—Study 1

Data from the Naturalistic Neuroimaging Database^50^ was used for this study. In this database, participants watched a range of full-length movies that contained rich social and emotional content. Their only task was to watch the movie, which makes this data ideal for investigating how signals of emotion might be encoded in naturalistic contexts.

#### Participants

20 subjects were recruited from the London area (mean age = 27.7, 50% female, 30% Black, Asian, or ethnic minority). Participants were screened out if they had previously seen the movie (*500 Days of Summer*), so that all participants were viewing it for the first time when they completed the study. In addition, all participants were right-handed and native English speakers. Participants were excluded if they had a history of claustrophobia, psychiatric or neurological illness, if they were taking medication, or if they had a hearing or uncorrected visual impairment.

#### Paradigm

Brain activity was measured using fMRI while subjects viewed the full-length movie *500 Days of Summer*. The movie was presented in two ∼50 minute segments. Participants were able to pause in the middle of a segment if they needed a break, after which the scan and the movie were resumed in a synchronized manner (See “Movie pausing” in Aliko et al., 2020). After the movie was complete, an anatomical scan was acquired.

#### MRI acquisition

Data was acquired on a 1.5 T Siemens MAGNETOM Avanto with a 32 channel head coil (Siemens Healthcare, Erlangen, Germany). For functional images during movie-viewing, a multiband EPI sequence was used (TR = 1 s, TE = 54.8 ms, flip angle of 75°, 40 interleaved slices, resolution = 3.2 mm isotropic). For the anatomical image, a 10-minute high-resolution T1-weighted MPRAGE was collected (R = 2.73 s, TE = 3.57 ms, 176 sagittal slices, resolution = 1.0 mm).

#### Preprocessing

Preprocessing was carried out through AFNI and included slice-time corrections, de-spiking, volume registration, alignment to the MNI template, 6 mm smoothing, detrending with 6 motion regressors, timing correction to account for pauses in scanning, concatenation of runs across the movie, and manual artifact removal with ICA (for a detailed description of all preprocessing steps, see “Preprocessing: Functional” in Aliko et al., 2020).

### fMRI Data—Study 2

Data collected by Noad and colleagues^63^ was used for Study 2. In this experiment, participants watched a series of short audiovisual clips taken from the television series *Game of Thrones*. As in Study 1, participants were instructed to passively view the clips without completing any concurrent task.

#### Participants

45 neurologically healthy participants (median age = 19, 66% female) were recruited to participate in the study at the York Neuroimaging Centre at the University of York. 23 participants were familiar with *Game of Thrones*, while the remaining 22 were unfamiliar with the show.

#### Paradigm

Brain activity was measured using fMRI while subjects viewed a set of short audiovisual clips that ranged from about 1-2 minutes in duration. In total, the stimulus was about 13 minutes long. In addition, T1 anatomical images were obtained, as well as category localizer scans for faces, scenes, and scrambled faces.

#### MRI Acquisition

Data were acquired on a 3 T Siemens MAGNETOM Prisma with a 64 channel head coil (Siemens Healthcare, Erlangen, Germany). For functional images during stimulus viewing, a multiband EPI sequence was used (TR = 2 s, TE = 30 ms, flip angle of 80°, 60 interleaved slices, 2 mm isotropic resolution). For the anatomical image, a high-resolution T1-weighted MPRAGE was collected (TR = 2.3 s, TE = 2.26 ms, resolution = 1.0 mm).

#### Preprocessing

Data were preprocessed using the standard fMRIPrep pipeline^83^, which includes slice-time correction using AFNI’s 3dTshift^84,85^ and motion correction using FSL’s mcflirt^86^. In addition, 6 mm smoothing was conducted using AFNI’s 3dBlurToFWHM^87^, the same procedure employed in Study 1.

### Analyses

#### Definition of Regions of Interest

Regions of interest representing the posterior STS and amygdala were obtained from cortical and subcortical atlases, respectively. For posterior STS, the dorsal and ventral portions of posterior STS (“STSdp” and “STSvp”) from a multimodal parcellation of the human cortex were selected^58^. For the FFA and OFA, corresponding regions from the same multimodal cortical parcellation were selected (“FFC” and “PIT”, respectively). For the amygdala, anatomical subdivisions based on cytoarchitecture^88^ were selected from the SPM anatomy toolbox (including superficial, basolateral, centromedial, and amygdalostriatal areas). For the analysis exploring lateralization effects, regions of interest were split by hemisphere.

#### Encoding Model Creation from ANNs

To test whether representations of facial expression category or visual emotion schema are encoded in brain activity in the amygdala and posterior STS, we used two ANNs to process the movie and generate encoding models. One, called EmoFAN, is a deep convolutional neural network trained to classify the emotional expression of faces^46^. EmoFAN uses a face-alignment network^89^ to identify facial landmarks, which are then applied through an attention mechanism to a set of four convolutional blocks (each containing three convolutional layers). The final layer of EmoFAN has 10 dimensions and contains probabilities that the current stimulus falls into 1 of 8 emotion categories (neutral, happy, sad, surprise, fear, disgust, anger, and contempt) as well as continuous values for valence and arousal. The other, named EmoNet, is a deep convolutional neural network trained to classify the emotion schema evoked by a frame of visual input—i.e., visual context. EmoNet is a convolutional neural network based on AlexNet that consists of five convolutional layers and three fully connected layers and was trained to identify the human-rated emotional state of images from emotional video clips (for emotion categories, see Kragel et al., 2019). To develop the encoding models, we fed every fifth frame of the movie into each ANN and extracted activations from the intermediate and late layers of each model. The intermediate layer for EmoFAN was the concatenated convolutional layers of the final convolutional block, while the intermediate layer for EmoNet was the penultimate fully connected layer; the late layers were the last fully connected layer for both ANNs.

#### Comparison with Brain Data

Having obtained features from both ANNs for each frame of the video, we next convolved these features with the hemodynamic response function to generate an expected response if a given brain region was encoding these features. Using these features, we used partial least squares (PLS) regression to search for a model that explained BOLD activity in the region of interest. For EmoFAN, a 10-dimensional PLS regression model was used; for EmoNet and the combined model, 20 dimensions were used. Using 5-fold cross-validation, we trained the model on four-fifths of the subjects’ data and tested on the remaining fifth by calculating the correlation between predicted and actual brain activity. Following other studies using encoding models to characterize sensory processing^90–93^, we used the prediction-outcome correlation as a measure of performance, as it represents information content that could be read out by downstream regions.

#### Noise Ceiling Calculations

To estimate the noise ceiling for each region of interest, we performed PLS regression using resubstitution. That is, we trained a PLS regression model on all of each subject’s data and tested it on the same data, which provides an upper bound for how much variance a model of a given complexity would ever be able to explain (for a complete list of noise ceiling estimates, see Supplementary Table 2). Noise ceiling estimates are indicated in the figures with a vertical line.

#### Representational Similarity Analysis

We conducted a representational similarity analysis to examine the relationship between encoding model performance in multiple regions of the face processing network. For each subject, we correlated predictions of the encoding models between pairs of regions (posterior STS, amygdala, FFA, OFA), and then compared the average correlation between different sets of regions across subjects using paired t-tests on Fisher transformed correlation coefficients.

#### Statistical Tests

First, we performed *t*-tests to determine whether the average prediction-outcome correlation was above zero. Next, we analyzed the model weights (β_PLS_) to determine which voxels were predicted by our encoding models across subjects. A false discovery rate correction of *q* = .05 was used as a threshold to determine significance. To compare the effects of model, region, and depth, we performed ANOVAs on the average prediction-outcome correlation across subjects. We performed two-way ANOVAs to examine the effect of model and region, and we did this for late and intermediate layers. In addition, we performed a three-way ANOVA to test the effects of model, region, and depth all together.

## Author Contributions

P.K. conceived of the study design; P.K., K.S., and G.J. analyzed the data; P.K. and K.S. interpreted the results; K.S. wrote the manuscript; P.K. and G.J. edited the manuscript.

## Code Availability

The code for this study is available at the following GitHub repository: https://github.com/ecco-laboratory/SEE

## Data Availability

Data from Study 1 are available from the open-source Naturalistic Neuroimaging Database (see Aliko et al. 2020). Data from Study 2 are available on OpenNeuro (ds004848, see Noad et al.). The data used to fine tune and evaluate EmoNet can be obtained upon request from the authors of Cowen & Keltner, 2017. The KDEF can be obtained upon request from https://kdef.se/download-2/

## Statement on Human Research Participants

Data from both studies complies all relevant ethical regulations; data analyzed in Study 1 was approved by the ethics committee of University College London; data analyzed in Study 2 was approved by the ethics committee of the York Neuroimaging Centre (University of York, York, UK). All participants provided informed consent.

## Supplementary Text

The use of encoding models to test claims about how different perceptual, cognitive, and affective variables are represented in neural activity is a mainstream approach in neuroscience^1,2^. Although it is common to use experimentally manipulated variables as inputs to an encoding model (with outputs corresponding to firing rates or patterns of BOLD response), in the last decade it has become mainstream to use variables (often referred to as “representations”) from artificial neural networks that have been trained to perform different tasks for this purpose^3,4^. The basic logic of this approach is that if an ANN is trained to accomplish a given task (e.g., object recognition), representations learned by the ANN will be sufficient to predict activity in brain systems involved in that task (e.g., the ventral visual stream), whereas representations from ANNs trained for other tasks will be less predictive. Thus, the performance of ANNs as feature extractors for encoding models can be used as a metric to make inferences about the types of representations encoded in different brain regions. In the present study, we used two deep convolutional neural networks that differ in their training data and objectives to model the activity of distributed face systems during free viewing of a cinematic film. The nature of representations learned by these networks differs, as EmoFAN was trained to classify emotional facial expressions from cropped images of faces^5^ and EmoNet was trained to classify the emotion schemas (based on the broader visual context) of frames taken from short video clips with varied content^6^. Because some of the videos used to train EmoNet include facial expressions, it is possible that information conveyed by the face could be used to classify certain emotion schemas (e.g., if ‘disgust’ videos contained facial expressions of retching following the consumption of noxious or spoiled food). That is, EmoNet may have learned representations of facial expression despite not being trained for this objective. Conversely, it is possible that information about facial expressions learned by EmoFAN can be used to classify emotion schemas, to the extent that those signals are present in the sensory array.

To evaluate the sensitivity of EmoFAN and EmoNet to variation in different sources of visual information, we probed representations in the last fully connected layer of each model using multivariate classification and 5-fold cross-validation in a face database (the Karolinska Directed Emotional Faces, KDEF)^7^ and an emotional scene database (see Materials and Methods for details). If each ANN learned representations that are highly specialized for a single task, then one would expect to observe a dissociation in classification accuracy such that EmoFAN was sensitive to changes in facial expression (but not context) and EmoNet would be sensitive to differences in general context (but not facial expression). Alternatively, if the representations learned for one objective are useful for multiple behaviors, then both classifiers should perform above chance, albeit with different performance levels and mappings between sensory features and emotion categories.

This sensitivity analyses revealed that features from the two ANNs differed in their ability to classify the emotional content of frames from short video clips (EmoFAN mean area under the receiver operating characteristic curve (AUC) = .700, *sem* = .018; EmoNet mean AUC = .847, *sem* = .019; Supplementary Figure 1). Classification performance was more comparable when using each model’s representations to classify expression categories from a dataset of emotional faces (EmoFAN mean AUC for KDEF = .658, *sem* = .013; EmoNet mean AUC for KDEF = .685, *sem* = .010, Supplementary Figure 2). These results suggest that information about emotion present in the face and the broader visual scene is present in late layers of both models, and that EmoNet is more sensitive to differences in the overall visual scene compared to facial expressions.

Inspecting how representations in late layers of ANNs were mapped onto categories of visual context and facial emotion revealed further differences between EmoFAN and EmoNet. Regression coefficients from visual scene classifiers showed that multiple EmoFAN categories were used to map facial expression information onto any scene category, while EmoNet’s mappings, on the whole, contain more one-to-one mappings. Regression coefficients from expression classifiers showed that sensible one-to-one mappings between EmoFAN’s representations and certain expression categories exist (e.g., fear, and surprise), whereas the mappings between EmoNet and facial emotions are distributed in a many-to-one manner, suggesting they could reflect statistical regularities in expression-context associations^8^ rather than individual facial expression categories.

Given the observed differences in the representations learned by EmoFAN and EmoNet in curated databases, we next evaluated whether they exhibit similarly distinct responses to the naturalistic stimuli used to develop fMRI encoding models in Study 1. To this end, we conducted a representational similarity analysis comparing the similarity of activations in intermediate and late layers of each ANN. This analysis showed that representations of the video stimulus used in Study 1 were strikingly different across ANNs, as the similarity of activation was higher within ANNs (*r* = .765, *sem* = 1.88 x 10^-5^) compared to between ANNs (*r* = -.092, *sem* = 4.25 x 10^-6^; Supplementary Figure 3). This demonstrates that EmoFAN and EmoNet capture unique information related to the face and visual scene, and that encoding models built using these representations could be mapped onto dissociable brain systems, if they exist.

**Supplementary Figure 1.**
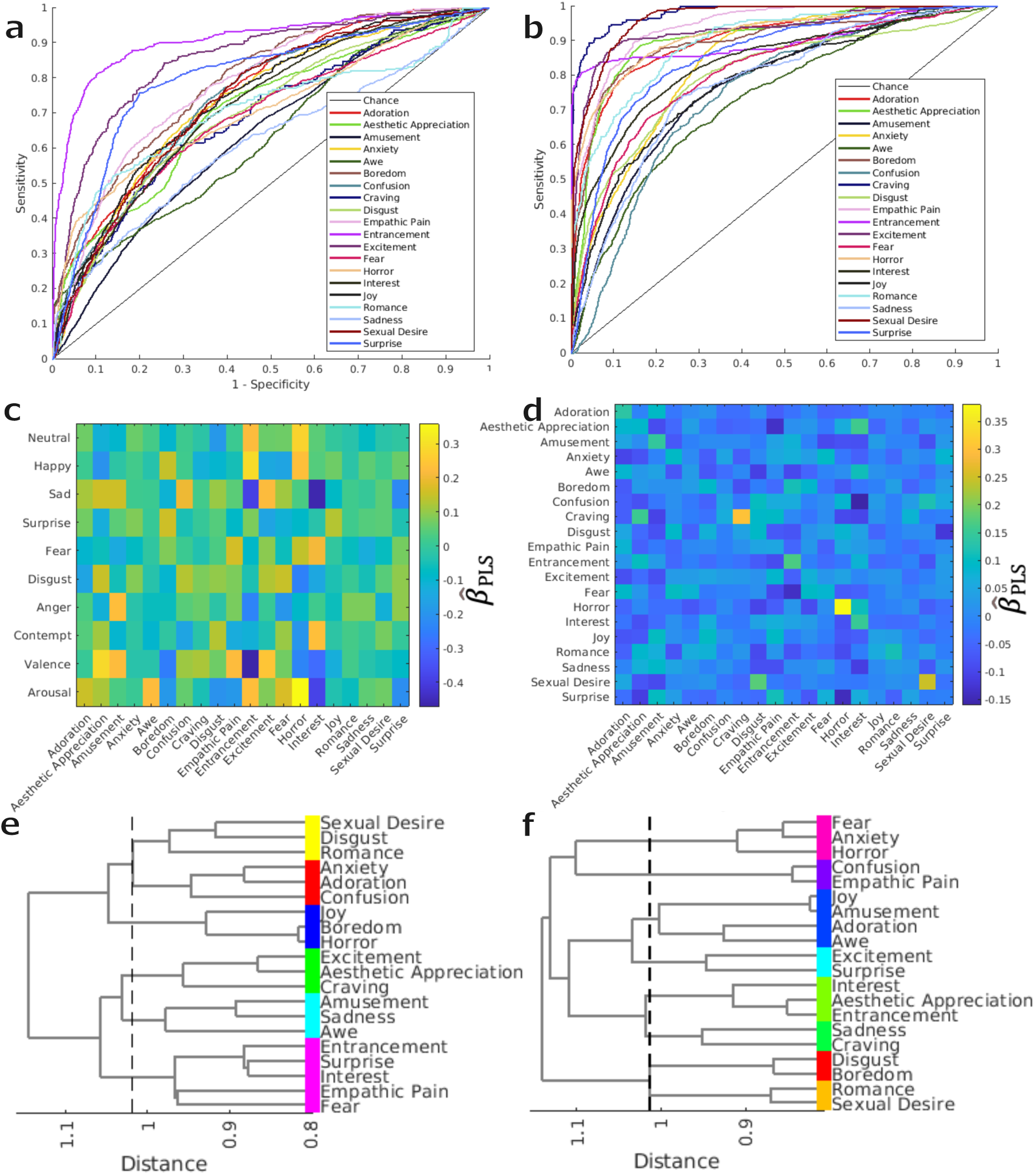
Classifier built using features from EmoNet (right) outperforms EmoFAN (left) at classifying the emotional context of frames from video clips in data collected by Cowen & Keltner. (**a** and **b**) Receiver operating characteristic curves showing classification performance for each category in the Cowen & Keltner dataset; chance indicated with solid black line (*N* = 24,632 image frames). (**c** and **d**) Betas from the regression model used to predict the emotion categories in the Cowen & Keltner dataset (x axis) based on the categories from the late layers of each model (y axis). (**e** and **f**) Results of hierarchical clustering performed on the confusion matrix for each classification. Dashed line indicates the cut point for the clustering solution in which each cluster is statistically distinguishable from the average of other clusters, with marginal colors indicating cluster assignment.

**Supplementary Figure 2.**
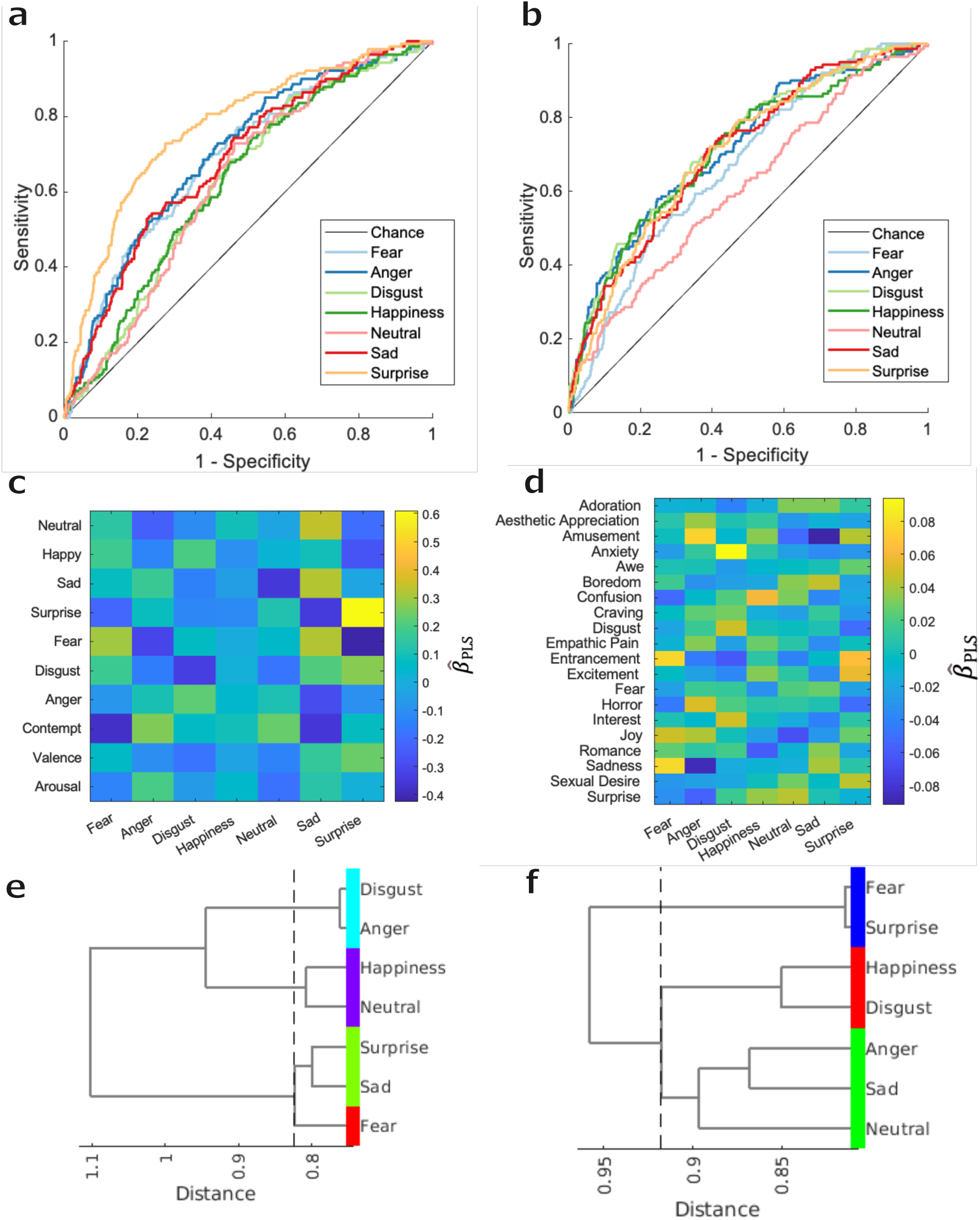
Classifier built using features from EmoNet (right) and EmoFAN (left) perform comparably at classifying the emotion category of faces from the Karolinska Directed Emotional Faces (KDEF) dataset using different mappings. (**a** and **b**) Receiver operating characteristic curves showing classification performance for each category in the KDEF dataset; chance indicated with solid black line (*N* = 981 images). (**c** and **d**) Betas from the regression model used to predict the emotion categories in the KDEF dataset (x axis) based on the categories from the late layers of each model (y axis). (**e** and **f**) Results of hierarchical clustering performed on the confusion matrix for each classification. Dashed line indicates the cut point for the clustering solution in which each cluster is statistically distinguishable from the average of other clusters, with marginal colors indicating cluster assignment.

**Supplementary Figure 3.**
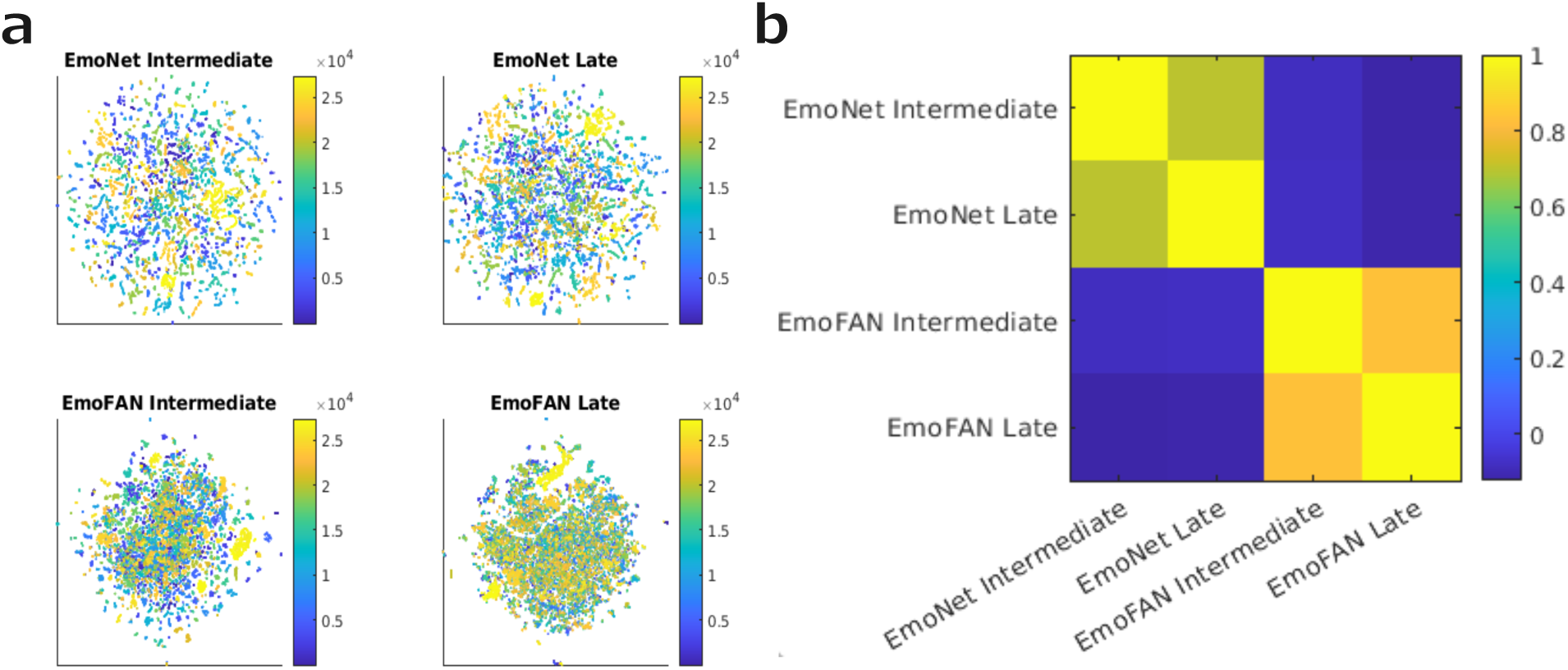
Features extracted from frames of the video stimulus in Study 1 (500 Days of Summer) by intermediate and late layers of EmoNet and EmoFAN are distinct. (a) t-distributed Stochastic Neighbor Embedding (*t*-*SNE*) plots showing features from the video over time (*N* = 136,777 frames). Color indicates the frame of the movie, with early frames represented in blue and later frames in yellow. Each *t-SNE* plot is initialized at the same points, so similar representations should produce similar plots. (b) Representational similarity matrix showing the Pearson correlation between video-derived features from EmoNet and EmoFAN. Yellow indicates a strong positive relationship.

**Supplementary Figure 4.**
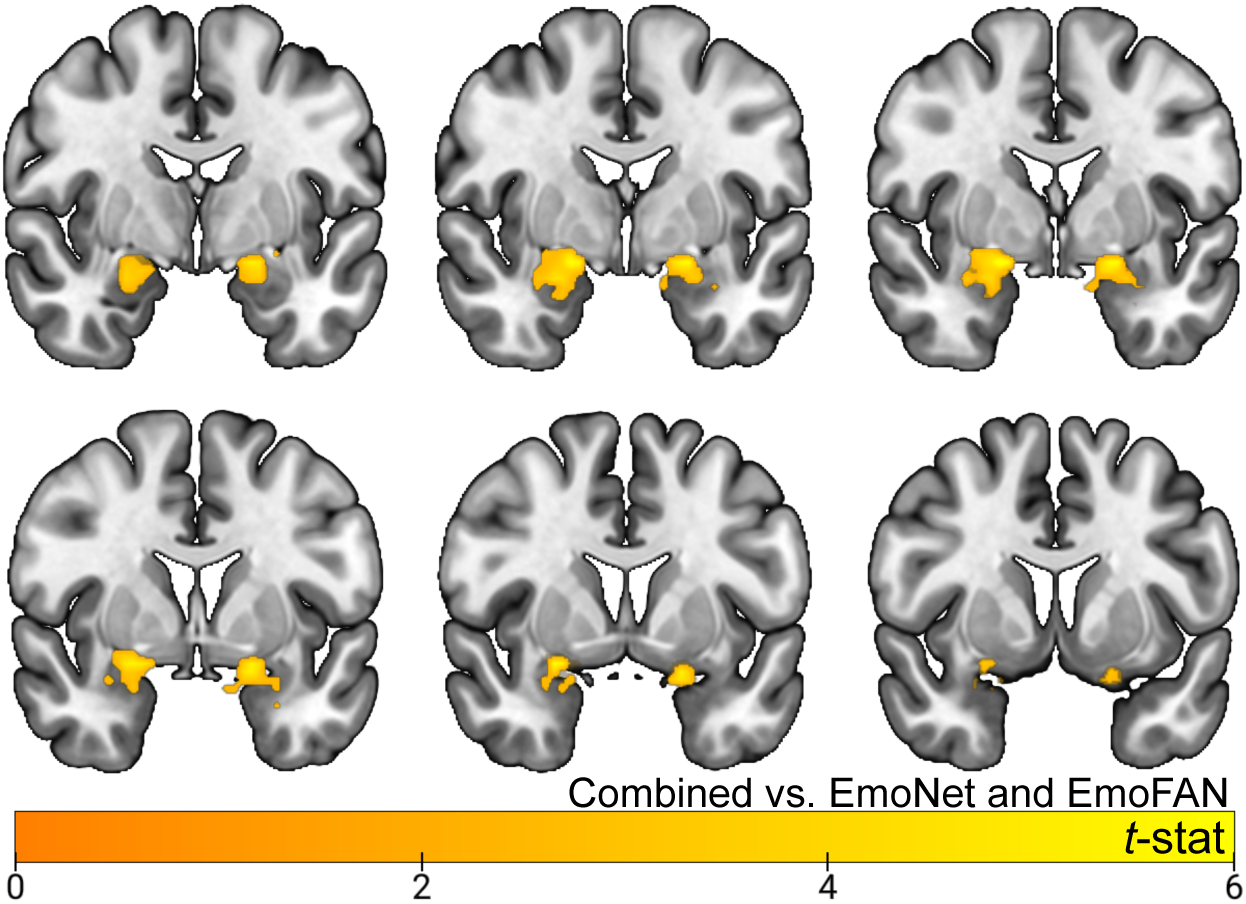
Encoding model performance for features from the late layers of each model predicting amygdala activity. The contrast shown here indicates which amygdala voxels are better predicted by the combined model compared to either the facial expression or context model alone, *q* < .05.

**Supplementary Figure 5.**
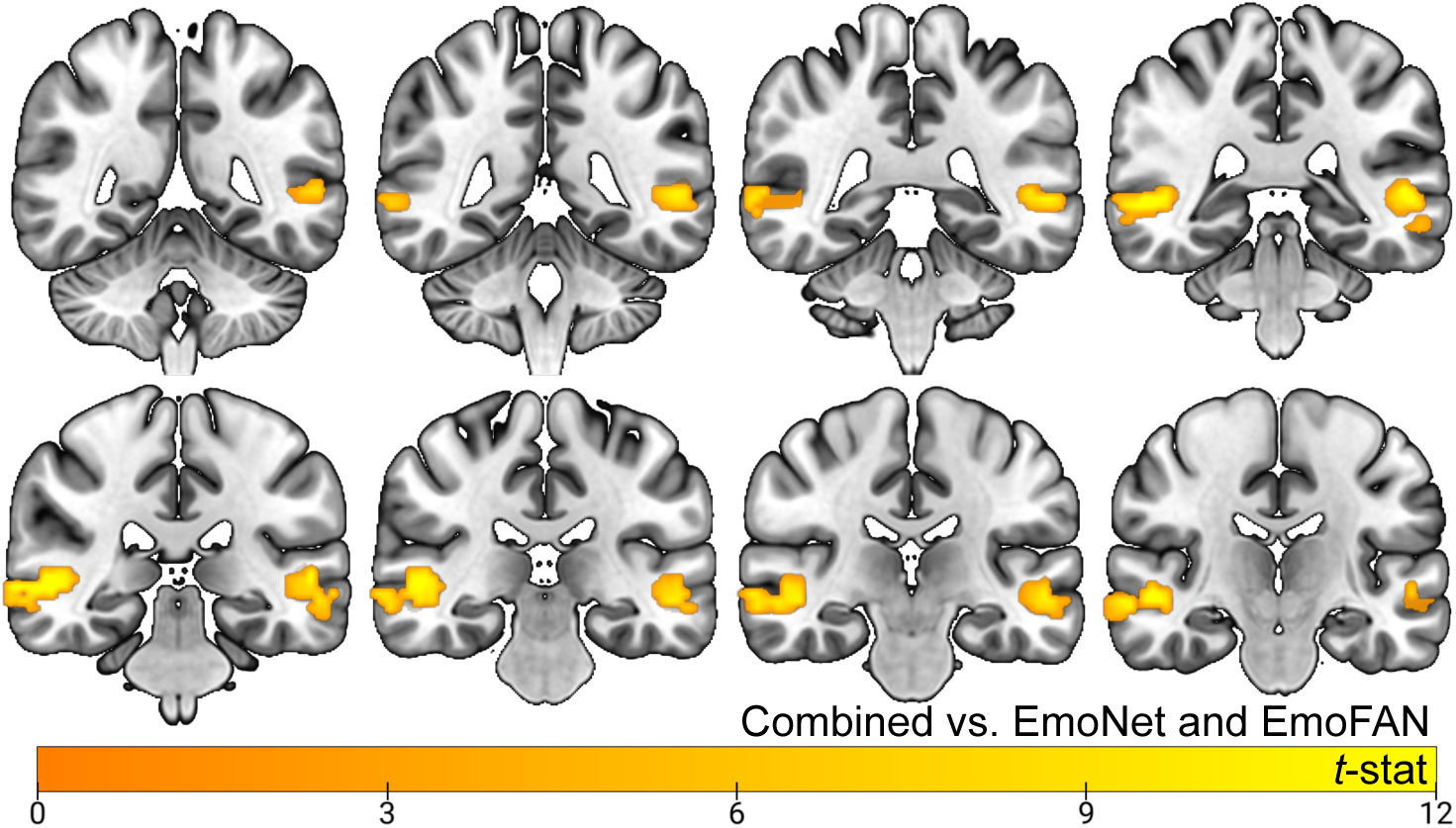
Encoding model performance for features from the late layers of each model predicting posterior STS. The contrast shown here indicates which STS voxels are better predicted by the combined model compared to either individual model, *q* < .05.

