## Supplemental Tables for "Sensory encoding of emotion conveyed by the face and visual context"

### Supplementary Tables

Supplementary Table 1. Raw prediction-outcome correlation in the amygdala and posterior STS

| | Amygdala | pSTS |
|---|---|---|
| EmoNet Late | *M* = .0094, *SD* = .0099, CI = (.0061, .0149) | *M* = .0864, *SD* = .0184, CI = (.0789, .0945) |
| EmoFAN Late | *M* = .0100, *SD* = .0071, CI = (.0072, .0135) | *M* = .0537, *SD* = .0121, CI = (.0495, .0600) |
| Combined Late | *M* = .0137, *SD* = .0110, CI = (.0098, .0194) | *M* = .0971, *SD* = .0194, CI = (.0891, .1057) |
| EmoNet Int | *M* = .0462, *SD* = .0224, CI = (.0385, .0584) | *M* = .2354, *SD* = .0423, CI = (.2178, .2535) |
| EmoFAN Int | *M* = .0368, *SD* = .0175, CI = (.0307, .0461) | *M* = .1715, *SD* = .0335, CI = (.1578, .1860) |
| Combined Int | *M* = .0634, *SD* = .0260, CI = (.0546, .0784) | *M* = .2624, *SD* = .0439, CI = (.2436, .2808) |

Supplementary Table 2. Mean percent noise ceiling (95% CI) in amygdala and posterior STS

| Features | Amygdala | pSTS |
|---|---|---|
| EmoNet Late | .1325 (.0873, .1951) | .7047 (.6735, .7310) |
| EmoFAN Late | .2005 (.1479, .2547) | .6780 (.6500, .7092) |
| Combined Late | .1793 (.1293, .2375) | .7190 (.6922, .7446) |
| EmoNet Int | .3595 (.3171, .4105) | .7999 (.7861, .8142) |
| EmoFAN Int | .4060 (.3615, .4616) | .8007 (.7840, .8174) |
| Combined Int | .4402 (.4049, .4902) | .8187 (.8055, .8315) |

Supplementary Table 3. Results of *t*-tests comparing late and intermediate layers of each model

| Features | Amygdala | pSTS |
|---|---|---|
| EmoFAN | $t = -9.17$, $p < .001$, 95% CI = (-.2525, -.1587) | $t = -11.04$, $p < .001$, 95% CI = (-.1459, -.0994) |
| EmoNet | $t = -10.74$, $p < .001$, 95% CI = (-.2712, -.1827) | $t = -8.75$, $p < .001$, 95% CI = (-.1179, -.0724) |

Supplementary Table 4. Mean prediction-outcome correlation for each model by hemisphere

| | Amy left | Amy right | pSTS left | pSTS right |
|---|---|---|---|---|
| EmoNet Late | .0106 | .0105 | .0853 | .0873 |
| EmoFAN Late | .0104 | .0110 | .0501 | .0568 |
| Combined Late | .0142 | .0146 | .0964 | .0985 |
| EmoNet Int | .0451 | .0458 | .2614 | .2501 |
| EmoFAN Int | .0346 | .0354 | .1978 | .1794 |
| Combined Int | .0609 | .0621 | .2895 | .2772 |

Supplementary Table 5. Results of paired *t*-tests comparing left vs. right hemispheres

| | Amygdala | pSTS |
|---|---|---|
| EmoNet Late | $t$ = -.1449, $p$ = .8863, 95% CI: (-.0255, .0222) | $t$ = -1.1243, $p$ = .2749, 95% CI: (-.0850, .0256) |
| EmoFAN Late | $t$ = -.7856, $p$ = .4418, 95% CI: (-.0431, .0196) | $t$ = -1.7516, $p$ = .0960, 95% CI: (-.1129, .0100) |
| Combined Late | $t$ = -.5950, $p$ = .5589, 95% CI: (-.0391, .0218) | $t$ = -1.0536, $p$ = .3053, 95% CI: (-.0795, .0263) |
| EmoNet Int | $t$ = -2.0734, $p$ = .0520, 95% CI: (-.0647, .0003) | $t$ = -0.6687, $p$ = .5117, 95% CI: (-.0433, .0223) |
| EmoFAN Int | $t$ = -2.3873, $p$ = .0275, 95% CI: (-.067, -.0044) | $t$ = .2974, $p$ = .7694, 95% CI: (-.027, .0359) |
| Combined Int | $t$ = -2.7594, $p$ = .0125, 95% CI: (-.0627, -.0086) | $t$ = -.2487, $p$ = .8062, 95% CI: (-.0299, .0236) |

Supplementary Table 6. Mean percent noise ceiling (95% CI) for encoding model performance in core face regions

| | FFA | OFA |
|---|---|---|
| EmoNet Late | .7146 (.6878, .7405) | .7457 (.7106, .7692) |
| EmoFAN Late | .6693 (.6285, .7028) | .6948 (.6469, .7319) |
| Combined Late | .7164 (.6918, .7407) | .7423 (.7157, .7650) |
| EmoNet Int | .8001 (.7790, .8146) | .8065 (.7887, .8235) |
| EmoFAN Int | .7849 (.7619, .8037) | .7832 (.7593, .8047) |
| Combined Int | .8034 (.7862, .8183) | .8018 (.7781, .8195) |

Supplementary Table 7. Raw prediction-outcome correlation (95% CI) in the amygdala and posterior STS for Study 2

| | Amygdala | pSTS |
|---|---|---|
| EmoNet Late | *M* = .3875, CI = (.3622, .4141) | *M* = .4005, CI = (.3818, .4185) |
| EmoFAN Late | *M* = .2511, CI = (.2303, .2736) | *M* = .2658, CI = (.2540, .2793) |
| Combined Late | *M* = .4246, CI = (.3944, .4516) | *M* = .4329, CI = (.4144, .4512) |
| EmoNet Int | *M* = .5478, CI = (.5134, .5801) | *M* = .5895, CI = (.5674, .6114) |
| EmoFAN Int | *M* = .4571, CI = (.4272, .4880) | *M* = .4734, CI = (.4536, .4918) |
| Combined Int | *M* = .5437, CI = (.5092, .5759) | *M* = .5860, CI = (.5629, .6070) |
